# African Green Monkey Cerebrospinal Fluid miRNome Captures Conserved miRNAs Relevant to Human Neurodegenerative Disease

**DOI:** 10.64898/2026.08.07.743104

**Authors:** Sara Dzigurski, Rashid Al-Abri, Xiaoting Li, Monica R. Grasty, Alice C. Rodrigues, Michael R. Weed, John D. Elsworth, Matthew S. Lawrence, Yujing J. Heng, Cristina S. Bogsan, Pourya Naderi Yeganeh, Winston A. Hide, Frank J. Slack, Gamze Gursoy, Andrew D. Miranker, Bianca R. P. Brown

## Abstract

**Background:** The African green monkey (AGM) is increasingly used as a model for early-stage Alzheimer’s disease (AD), with cerebrospinal fluid (CSF) targeted for biomarker discovery and longitudinal disease monitoring of shifts in the central nervous system. MicroRNAs (miRNAs) are particularly informative indicators of early neuropathological change. Despite the complementary value of an early-stage disease model and a molecular marker capable of capturing early change, the miRNA composition (miRNome) of AGM remains undefined. We established the AGM CSF miRNome from antemortem samples using miRNA sequencing and a qRT-PCR-based array. We also developed a hierarchical annotation pipeline to classify miRNAs as either family-conserved or unclassified and to assess sequence alignment across humans and other species.

**Results:** We used untargeted miRNA sequencing to characterize the AGM CSF miRNome and identified 205 miRNAs that could be classified into three family-conserved categories: canonical, noncanonical, and 3′-terminal variants. Of these, 150 were also detected using a human-targeted qRT-PCR array, providing independent support for the sequence-derived miRNome. Sequencing abundance and qRT-PCR array Ct values showed significant cross-platform concordance overall, although concordance was lower for 3′-terminal isomiRs than for canonical miRNAs. Comparison with human GTEx tissue-expression data indicated that several human homologs of AGM CSF miRNAs exhibited brain-preferential expression. Notably, predicted targets of many of these miRNAs were enriched for pathways implicated in neurodegenerative disease. Finally, we identified 20 unclassified candidates that could not be assigned to established miRNA families, two of which we propose as putatively novel miRNAs.

**Conclusion:** The AGM CSF miRNome is substantially conserved with the human miRNome but also contains 3′-terminal isomiRs and unclassified miRNA candidates. AGM CSF contains miRNAs homologous to human miRNAs associated with AD and other neuropathologies, highlighting the translational potential of this model. However, our study also reveals challenges related to species-specific sequence variation and reduced cross-platform concordance for isomiRs. Thus, comparative studies will be needed to validate the functional and biomarker relevance of these miRNAs across species. More generally, this initial miRNome provides a reference resource for future studies of miRNAs in AGM across disease-related, physiological, experimental, and evolutionary contexts.

## INTRODUCTION

The African green monkey (AGM; *Chlorocebus sabaeus*) is emerging as a model for early neuropathological biomarker discovery in Alzheimer’s disease (AD). As they age, AGMs naturally accumulate amyloid and tau pathology without overt cognitive decline, making them particularly well suited for studying early-stage AD^1–3^. Unlike AGMs, standard rodent models do not spontaneously reproduce this combination of AD-like features without genetic or experimental manipulation^4^. Moreover, the closer evolutionary relationship between nonhuman primates and humans provides an additional biological basis for evaluating the translational relevance of this model. Recent work has shown that Aβ oligomer inoculation induces early AD-like molecular changes detectable in the CSF proteome, supporting the use of this biofluid as a less invasive source of brain-associated molecular information^3^. A key advantage of expanding the molecular toolkit for CSF analysis in an experimentally induced AGM model of AD is the ability to collect samples longitudinally. This enables molecular changes to be tracked before and during the earliest stages of disease progression, a period that is extremely difficult to capture prospectively in humans.

MicroRNAs (miRNAs), which act as post-transcriptional regulators of gene expression, are particularly informative indicators of early neuropathological change. These small noncoding RNAs regulate gene expression in eukaryotes through interactions with target mRNAs^5^. Despite the low RNA content of CSF, miRNAs are among the most consistently detectable RNA species in this fluid, making them well suited for CSF-based biomarker studies^6^. In AD, miRNAs have been implicated in synaptic dysfunction, Aβ and tau regulation, and neuroinflammation^7^. More than 30 miRNAs have also been proposed as potential CSF biomarkers of AD^6,7^. Although miRNAs are broadly conserved across primates^8^, the AGM miRNome remains poorly defined; therefore, the composition and abundance of miRNAs detectable in AGM CSF are largely unknown. Cross-species variation in mature sequence, processing, and annotation can further complicate miRNA identification and detection^9^. These gaps limit our ability to determine how closely AGM miRNAs resemble known human miRNAs, which miRNAs can be reliably recovered from CSF, and whether the CSF miRNome includes established miRNA biomarkers.

Two primary approaches are used for high-throughput miRNA quantification. Next-generation sequencing (NGS) provides an untargeted survey of miRNAs and enables detection of previously unannotated or novel miRNAs, but it generally requires higher RNA concentrations^10–12^. In contrast, qRT-PCR array-based platforms, while more sensitive, quantify a defined panel of known miRNAs^6,7^. Although arrays are widely used in biomarker studies, they depend on well-characterized miRNAs to build probes and are designed around human annotations. Human miRNA qRT-PCR array platforms have been applied to blood samples in nonhuman primates^13^ and tissue from other species^14^. As such, qRT-PCR can provide complementary information in unannotated species, with sequencing enabling broad discovery and sequence-level characterization and qRT-PCR providing sensitive detection of predefined miRNAs.

Here, we characterize the AGM CSF miRNome using antemortem CSF samples collected from six experimentally naïve females. We performed untargeted miRNA sequencing and developed a hierarchical annotation pipeline that integrates family- and sequence-level assignments to define the AGM CSF miRNome. We then applied a human-targeted qRT-PCR array as an independent platform to provide orthogonal support for the subset of human-annotated miRNAs detected by both miRNA-seq and the array. Finally, for miRNAs conserved between AGMs and humans, we integrated target prediction, tissue-expression, and enrichment analyses to identify putative regulatory functions and disease-associated pathways. More broadly, this study addresses the challenge of defining a biofluid miRNome in a translational species without a dedicated annotated miRNA reference and evaluates the constraints of cross-species miRNA profiling.

## RESULTS

### Study Overview

We characterized the CSF miRNome from six experimentally naïve female AGMs aged 7-9 years. CSF was collected from living individuals via a cisterna magna tap (up to 1 mL per tap). Total RNA was extracted from 100 µL of CSF, and concentrations ranged from 15-84 pg/µL. We first used miRDeep2^15^ for genome-based miRNA prediction to identify provisional candidate sequences from miRNA-seq data and assign provisional IDs (Figure 1; Table S1). We developed a hierarchical annotation pipeline to assign provisional miRNA candidates to annotated miRNAs. Our pipeline resolved family-level conservation using MirMachine^16^ and Rfam^17^, then we refined sequence-level identity by aligning to known miRNAs in the human-only and pan-species MirGeneDB databases^18^ (see Methods). We then performed targeted quantification using RNA extracts from the same CSF samples with the qRT-PCR-based PanoramiR panel^19^, which includes 376 human biofluid miRNA primers (Table S2). This provided independent detection to support the sequencing derived AGM CSF miRNome (Figure 1). We identified miRNAs detected by both miRNA-seq and qRT-PCR array and quantified cross-platform concordance. miRNA tissue-expression patterns were assessed using the gene expression quantifications from 54 tissues and 946 human postmortem donors of Genotype-Tissue Expression (GTEx) project^20^. Finally, to evaluate potential functions, we performed mRNA target prediction for miRNAs with known human annotations using miRWalk^21^. Target genes were then analyzed for functional and disease enrichment using Gene Ontology and Disease Ontology datasets.

**Figure 1.**
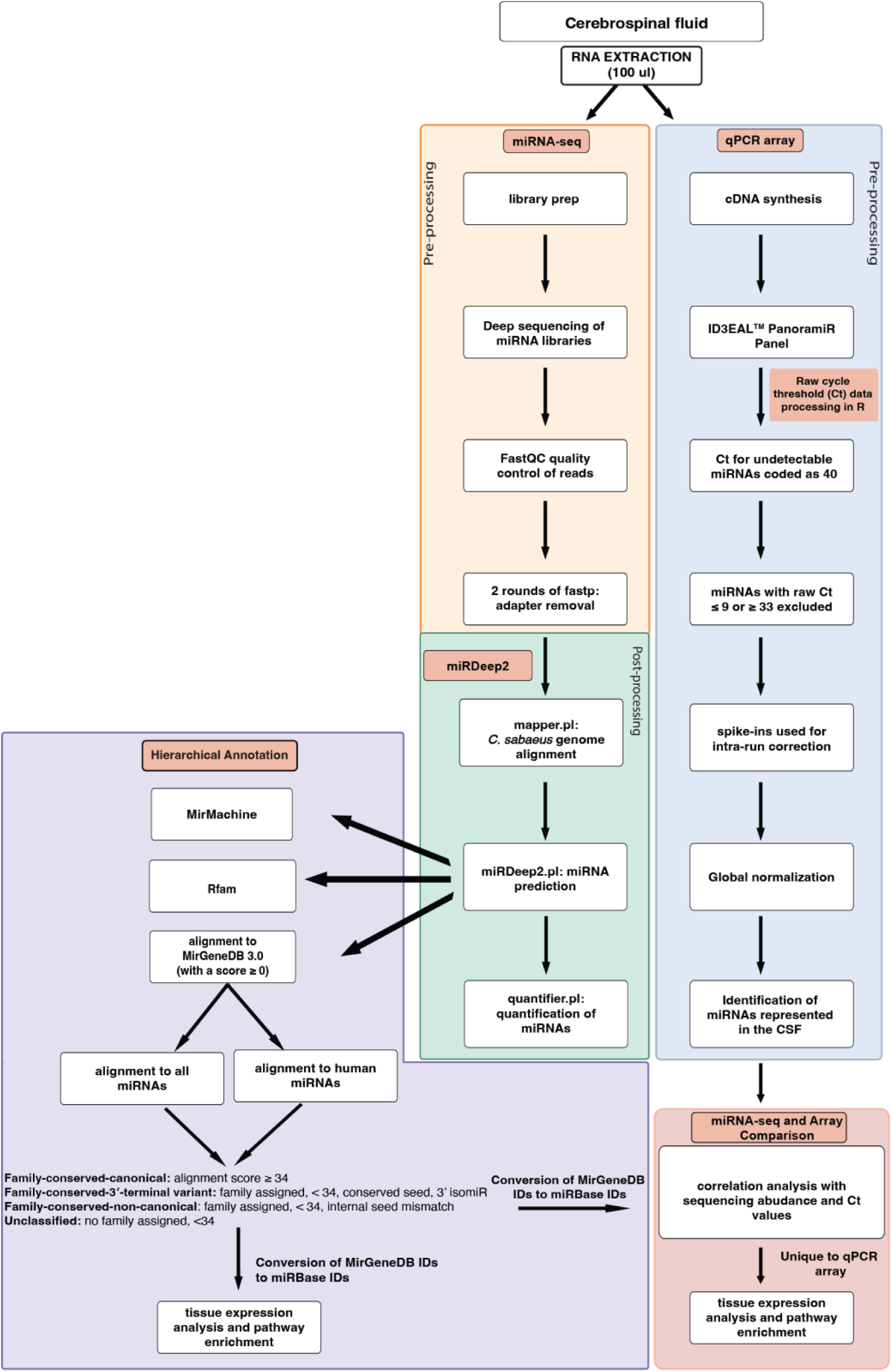
Experimental and analytic workflow to characterize African green monkey CSF miRNome. RNA extraction: RNA was extracted from 100 µL of CSF, and aliquots were designated for either miRNA sequencing or qRT-PCR array analysis. **miRNA-seq:** Small RNA libraries were prepared and sequenced. **miRDeep2:** Preprocessed reads were used as input for miRDeep2 for miRNA prediction and quantification. **Hierarchical annotation:** The resulting miRNA candidates were processed through our hierarchical annotation pipeline and classified as family-conserved canonical, family-conserved 3′-terminal variant, family-conserved noncanonical, or unclassified. **qRT-PCR array:** cDNA was synthesized from the extracted RNA and analyzed using the ID3EAL™ PanoramiR Panel. **miRNA-seq and array correlation:** miRNA abundance estimates from sequencing were compared with Ct values from the PanoramiR array using correlation analysis.

### Family-conserved miRNAs showed strong sequence conservation with human

We identified 248 provisional miRNA candidates using miRDeep2. Provisional candidates were retained if they had a total read count ≥ 10 across all samples and a miRDeep2 score ≥ 5 (Figure S1A-B). The miRDeep2 score reflects confidence that a sequence conforms to a canonical miRNA structure. We developed a hierarchical annotation framework to classify miRDeep2 provisional miRNAs into four categories: family-conserved -canonical, -3′-terminal variant, -noncanonical and unclassified. First, we assigned provisional candidates to known miRNA families using MirMachine and Rfam. We then aligned each candidate separately against mature miRNA sequences in the human-only and pan-species MirGeneDB databases. The separate comparisons distinguished candidates matching human miRNAs from those matching miRNAs identified only in other species. Candidates with complete mature-sequence identity to an established MirGeneDB reference were classified as family-conserved canonical; all candidates in this category had alignment scores ≥34. In the benchmarking analysis, this threshold corresponded to fewer than two mismatches, no more than one internal indel, and no more than five indels at the 3′ terminus and yielded 99.7% specificity and 40.0% sensitivity (Figure S1D). Among candidates with alignment scores <34, those with family-level support from MirMachine or Rfam, identical 5′ and seed sequences to the assigned reference, no internal sequence differences, and variation confined to the 3′ terminus were classified as family-conserved 3′-terminal variants. Candidates with family-level support but with seed or internal sequence differences were classified as family-conserved noncanonical miRNAs. Candidates with alignment scores <34 and no family-level support were classified as unclassified and evaluated further as putatively novel miRNAs

Most detected AGM CSF miRNAs were family-conserved canonical and were annotated as human miRNAs. Across the 248 provisional miRNA candidates, mean reads per million (RPM) values varied widely with a mean of 2369.1 RPM (range, 0.05-234,000 RPM). The 248 provisional candidates corresponded to 225 unique miRNAs, comprising 205 family-conserved miRNAs and 20 unclassified candidates (Table S1). Among these 205 assigned miRNAs, 178 were classified as family-conserved canonical based on complete mature-sequence identity to known miRNAs, 24 as family-conserved 3′-terminal variants, and three as family-conserved noncanonical. Of the 205 miRNAs, 202 were assigned to a human reference.

The most abundant family-conserved miRNAs belonged to several highly conserved miRNA families. Among the family-conserved canonical miRNAs, Hsa-Mir-15-P2a_5p had the highest mean abundance (234,000 RPM, range = 93,200-338,000) and exceeded the next most abundant miRNA by approximately 6-fold. Other abundant miRNAs included Hsa-Mir-486_5p (40,400 RPM, range = 272-74,300), Hsa-Mir-26_5p (27,200 RPM, range = 1,350-61,400), and members of the let-7 family, including Hsa-Let-7-P2b2_5p (26,700 RPM, range = 15,500-44,800), Hsa-Let-7-P2c3_5p (13,900 RPM, range = 4,540-23,000), and Hsa-Let-7-P2c2_5p (13,800 RPM, range = 5,030-25,700). Additional abundant miRNAs included Hsa-Mir-204-P2_5p (9,290 RPM, range = 1,350-20,100), Hsa-Mir-17-P4d_5p (9,270 RPM, range = 5,500-13,600), Hsa-Mir-30-P1d_5p (8,000 RPM, range = 1,530-15,300), and Hsa-Mir-143_3p (7,800 RPM, range = 724-23,400). One additional family-conserved canonical miRNA aligned only to a nonhuman primate reference, Cja-Mir-193-P1a_5p, and had a mean abundance of 13.9 RPM (range = 0-76.5); it was detected in three samples.

We identified 24 family-conserved 3′-terminal variant miRNAs that were assigned to a family by Rfam and/or MirMachine but had an alignment score <34 (Figure 2B; Table S1). The low-alignment miRNAs were consistent with 3′ isomiRs, retaining identical 5′ and seed sequences with no internal sequence differences. These showed 2-5 nucleotide truncations at the 3′ terminus relative to the reference sequence. Among the top family-conserved 3′-terminal variants, the most abundant included members of the deeply conserved Let-7 and Mir-124 families. Hsa-Let-7-5p/Let-71-5p/Let-72-5p had the highest mean RPM (30,621.1 RPM, range = 8,699.0-71,203.2), followed by Hsa-Let-73-5p/Let-71-5p (10,407.0 RPM, range = 3,172.6-15,472.3), Hsa-Mir-124-v1-3p (2,727.0 RPM, range = 44.7-8,955.5), and Hsa-Let-7-P1b-5p (1,957.2 RPM, range = 350.0-4,611.4). In addition to the miRNAs assigned to human references, one family-conserved 3′-terminal variant aligned most closely to the macaque reference Mml-Mir-550-P1_5p.

**Figure 2.**
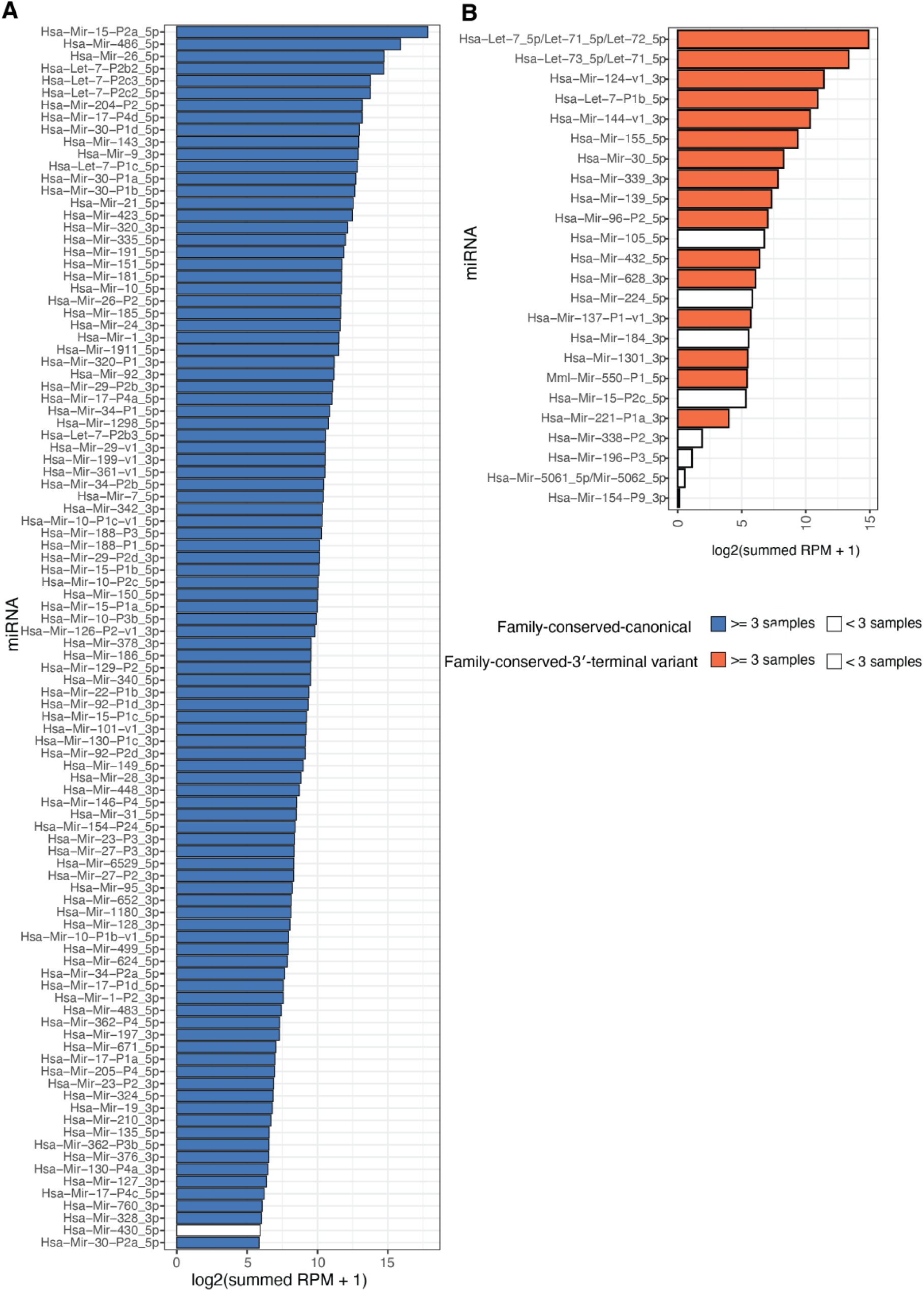
miRNAs identified in the African green monkey CSF miRNome. (A) Bar plot showing RPM for family-conserved canonical miRNA, defined by complete mature-sequence identity to an established MirGeneDB reference and have alignment scores ≥34. (B) Bar plot showing RPM for family-conserved 3′-terminal variant miRNA matches with family assignments supported by Rfam and/or MirMachine but alignment scores <34 relative to the best-matched reference miRNA. In both panels, the human miRNA ID is shown when a human match was identified; where no human match was identified, the best-matched nonhuman miRNA ID is shown. Filled bars represent miRNAs detected in ≥3 samples, whereas open bars represent miRNAs detected in <3 samples. The x-axis shows the log-transformed mean reads per million (RPM) for each miRNA.

We identified three family-conserved noncanonical miRNA candidates that had family-level support but alignment scores <34 and sequence variation within the seed region (Table S1). CM001961.2_13881 had the highest mean abundance (132.8 RPM, range = 0-342.2) and was assigned to the Mir-942 family by Rfam. CM001949.2_6595/CM001949.2_6206 (1.2 RPM, range = 0-4.1) were assigned to the mir-3158 family by Rfam, while CM001960.2_13216 (0.5 RPM, range = 0-2.8) was assigned to the Mir-991 family by MirMachine.

### Concordance Between miRNAs from miRNA-seq and the Array

We combined untargeted sequencing with the higher-sensitivity qRT-PCR array to provide orthogonal support to miRNA-seq defined miRNome, assess concordance between approaches, and evaluate the utility and limitations of applying an existing human-targeted platform to AGM CSF. Ct values from the array were globally normalized using the PanoramiR pipeline (see Methods). Of the 202 family-conserved miRNAs assigned to human (Hsa-) by MirGeneDB, 193 could be cross-referenced to miRBase. Of these 193, 150 (137 family-conserved canonical and 13 family-conserved 3′-terminal variant) were also detected by the qRT-PCR array. The remaining 43 miRBase annotated miRNAs from miRNA-seq data were not represented on the array panel.

Several of the miRNAs detected by miRNA-seq and array have been implicated in AD, including miR-16-5p, let-7i-5p, and miR-204-5p. In addition to the ten most abundant miRNAs, other family-conserved canonical miRNAs previously associated with AD-related pathways were detected, including miR-19b-3p, miR-29c-3p, miR-125b-5p, miR-15a-5p, miR-146a-5p, miR-206, miR-92a-3p, and miR-29b-3p. Notably, miR-16-5p is highly abundant in erythrocytes.

Among the 150 miRNAs detected by both miRNA-seq and the qRT-PCR array, 69 had Ct ≤ 25, 56 had 25 < Ct ≤ 30, and 25 had Ct > 30 (Figure 3B; Table S2). miRNAs detected by both miRNA-seq and the array had significantly lower Ct values than those detected only by the array (Wilcoxon test, P < 0.001; Figure 3C). Because lower Ct values correspond to higher miRNA abundance, this distribution (with most overlapping miRNAs falling at Ct ≤ 30) indicates that our AGM sequencing data is enriched for high-abundance miRNAs.

**Figure 3.**
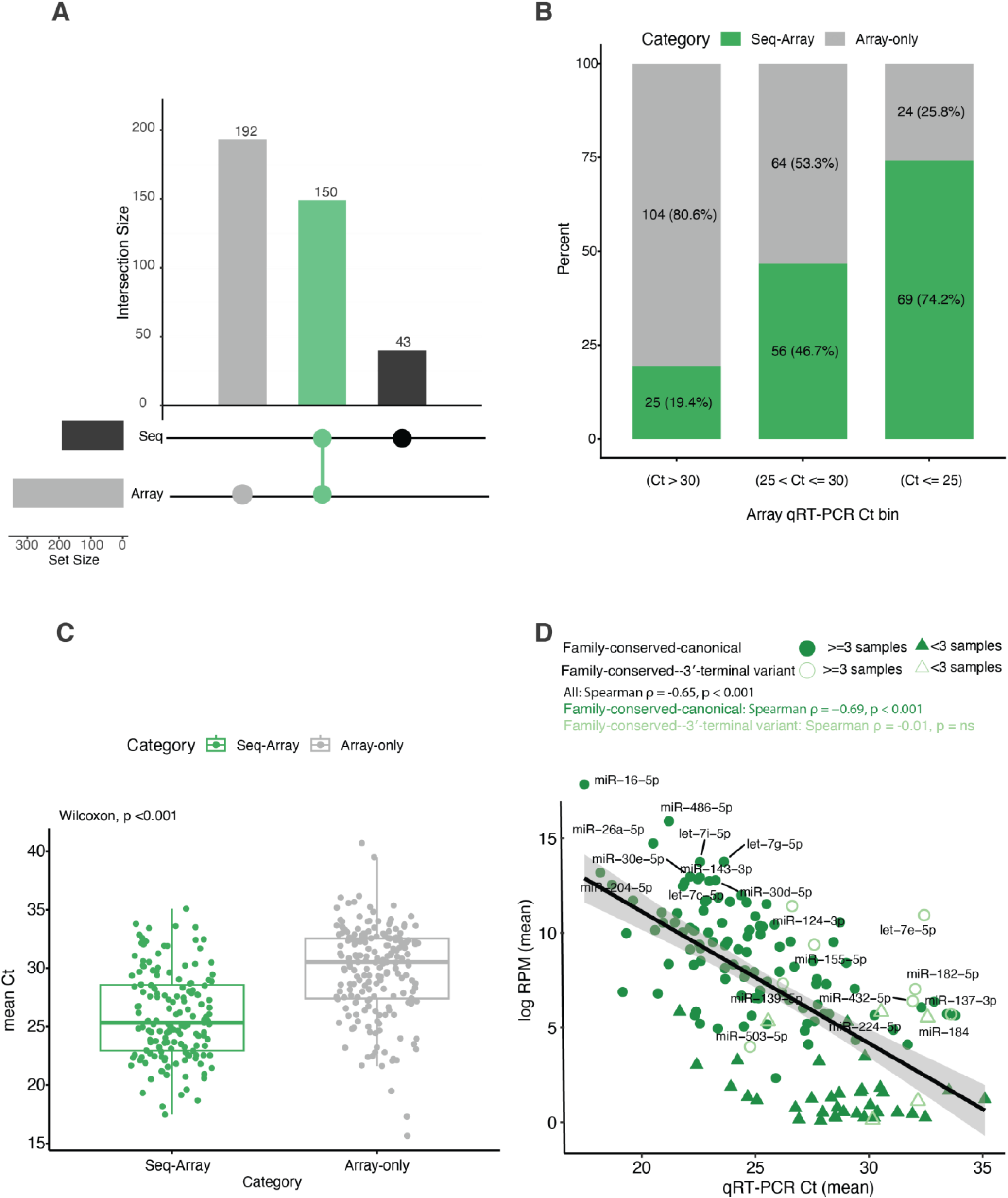
African green monkey CSF microRNAs identified by the miRNA-seq and qRT-PCR array methods. (A) Bar plot showing the number of unique and overlapping miRNAs detected by the array and sequencing platforms. The green bar indicates miRNAs detected by both platforms. (B) Proportion of miRNAs detected in the array dataset within each Ct category. MiRNAs detected by both miRNA sequencing and the qRT-PCR array are shown in green, whereas miRNAs detected only by the array are shown in gray. (C) Boxplot showing differences in mean Ct values of miRNAs detected by both platforms versus detected only by the array. (D) Correlation between array Ct values and mean sequencing abundance (RPM) for miRNAs detected by both methods. The 10 most abundant miRNAs in each of the family-conserved canonical and family-conserved 3′-terminal variant categories are labeled. Closed symbols indicate family-conserved canonical miRNAs, whereas open symbols indicate family-conserved 3′-terminal variant miRNAs. Circles indicate miRNAs detected in ≥3 individuals, and triangles indicate miRNAs detected in <3 individuals.

To assess concordance between platforms, we compared miRNA-seq sequencing read abundance with qRT-PCR normalized Ct values for overlapping miRNAs. Ct values and sequence abundance were significantly correlated across all miRNAs (Spearman ρ = −0.65, P < 0.001), indicating overall agreement between the two methods (Figure 3D; Table S2). This concordance was driven by family-conserved canonical miRNAs (Spearman ρ = −0.69, P < 0.001), because family-conserved 3′-terminal variants showed no significant correlation (Spearman ρ = −0.01, P = 0.99; Figure 3D; Table S2).

### Human homologs of AGM CSF miRNAs show preferential expression in brain tissues

To determine whether family-conserved canonical miRNAs detected in AGM CSF and miRNAs detected only by the qRT-PCR array showed tissue-preferential expression in humans, we compared their expression profiles across GTEx tissues. For the family-conserved canonical miRNAs, we identified a distinct cluster of 34 corresponding human miRNAs with preferential expression across multiple brain regions (Figure 4A-B; Table S3). Strong enrichment in the brain cortex was observed for Hsa-Mir-149_5p, Hsa-Mir-129-P2_5p, Hsa-Mir-128-P2_3p, Hsa-Mir-760_3p, Hsa-Mir-328_3p, and Hsa-Mir-331_3p. In the hypothalamus, Hsa-Mir-1298_5p, Hsa-Mir-448_3p, Hsa-Mir-138-P2_5p, and Hsa-Mir-154-P30_3p were enriched, while the nucleus accumbens (basal ganglia) showed enrichment for Hsa-Mir-1298_5p, Hsa-Mir-448_3p, and Hsa-Mir-592_5p. In addition to this brain-enriched cluster, we also observed miRNAs with high expression in non-brain tissues such as testis, blood, and kidney.

**Figure 4.**
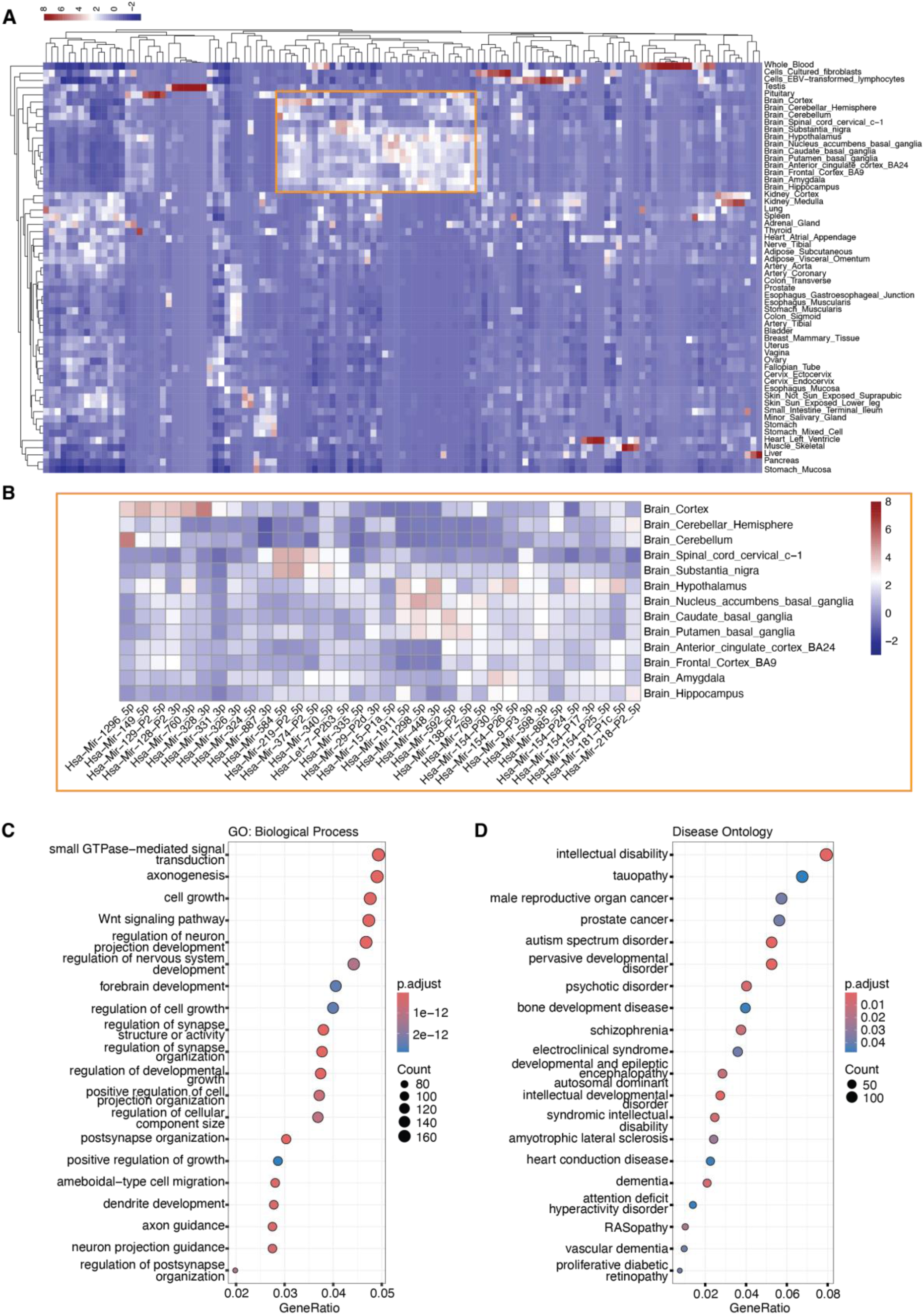
Tissue expression analysis of miRNAs and gene ontology (GO) and disease ontology (DO) enrichment of mRNA targets of African green monkey CSF miRNome. (A) Tissue-expression profiles of human homologs corresponding to family-conserved canonical AGM CSF miRNAs (alignment score ≥34 to a human reference and family support from MirMachine and/or Rfam), using GTEx v10, with (B) a zoomed-in view of the brain-tissue cluster. The top 20 enriched GO (C) or DO (D) terms for mRNAs predicted to be targets of these family-conserved canonical miRNAs by miRWalk, miRDB, and TargetScan. The GeneRatio indicates the number of mRNA targets to the total number of mRNA genes in the input list, while the terms are ranked by adjusted p-values.

Several array-unique miRNA showed preferential expression in brain tissues and were not detected by miRNA-seq. To determine whether miRNA-seq may have failed to detect brain-associated miRNAs detectable by the qRT-PCR array, we investigated the tissue expression patterns of miRNAs unique to the array. The array-unique set included 192 miRNAs (Figure S3). A subset of these demonstrated particularly strong brain expression, with heatmap intensities approximately two-fold or greater, including hsa-miR-9-5p, hsa-miR-329-3p, hsa-miR-410-3p, hsa-miR-431-3p, hsa-miR-433-3p, hsa-miR-107, hsa-miR-134-5p, and hsa-miR-212-3p (Figure S3).

### Predicted gene targets of family-conserved canonical miRNAs are enriched for neural pathways, neurodegenerative disease, and cancer

To assess whether predicted targets of the AGM CSF miRNome were associated with central nervous system processes, we performed Gene Ontology (GO) and Disease Ontology (DO) enrichment analysis on predicted mRNA targets identified using miRWalk for family-conserved canonical miRNAs with human assignments (Table S4-S5). GO enrichment revealed biological processes related to neuronal wiring and synaptic function, including axon guidance, dendrite development, and synapse organization. Predicted target genes were enriched for cellular-component terms including the synapse, dendritic spine, growth cone, and postsynaptic density (Figure S2B; Table S4-S5).

Consistent with these findings, Disease Ontology terms spanned neurodegenerative conditions (tauopathy, ALS, dementia, vascular dementia) and neurodevelopmental disorders (intellectual disability, autism spectrum disorder, ADHD; Figure 4D; Table S5). Targets were also enriched for cancer-related categories and epilepsy-related conditions (electroclinical syndrome and developmental and epileptic encephalopathy). These associations likely reflect shared regulatory pathways governing cell proliferation, differentiation, and apoptosis.

### Unclassified miRNAs in the AGM CSF miRNome

We identified 20 miRNAs that could not be assigned to a predicted gene family by MirMachine or Rfam and had low alignment scores when compared to known miRNAs. We therefore categorized these as unclassified (Figure 5; Table S1). None met our criteria for a close match to any mature miRNA in MirGeneDB across species, defined as fewer than two mismatches, no more than one internal indel, and no more than five terminal 3′ indels. Their best-matched reference miRNAs had alignment scores of 4-10, reflecting substantial sequence discordance due to mismatches and indels. All 20 miRNAs had total read counts >10 across samples, with mean RPM ranging from 0.05 to 149 RPM (Figure 5A; Table S1). Collectively, they accounted for <1% of total sequencing reads. Two were detected in four individuals and had relatively high miRDeep2 scores: CM001945.1_3281 (miRDeep2 score, 371.5; Figure 5B) and CM001963.2_15794 (miRDeep2 score, 246.9; Figure 5C). To further assess these candidates, their miRDeep2-predicted precursor sequences were queried against the NCBI nucleotide database using BLASTN^22^. The CM001945.1_3281 precursor showed 100% sequence identity to the putatively novel *Macaca fascicularis* miRNA precursor novel_miR60_3p (GenBank accession LR778035.1), suggesting that this sequence may be shared among nonhuman primates. In contrast, the CM001963.2_15794 precursor had no corresponding match.

**Figure 5.**
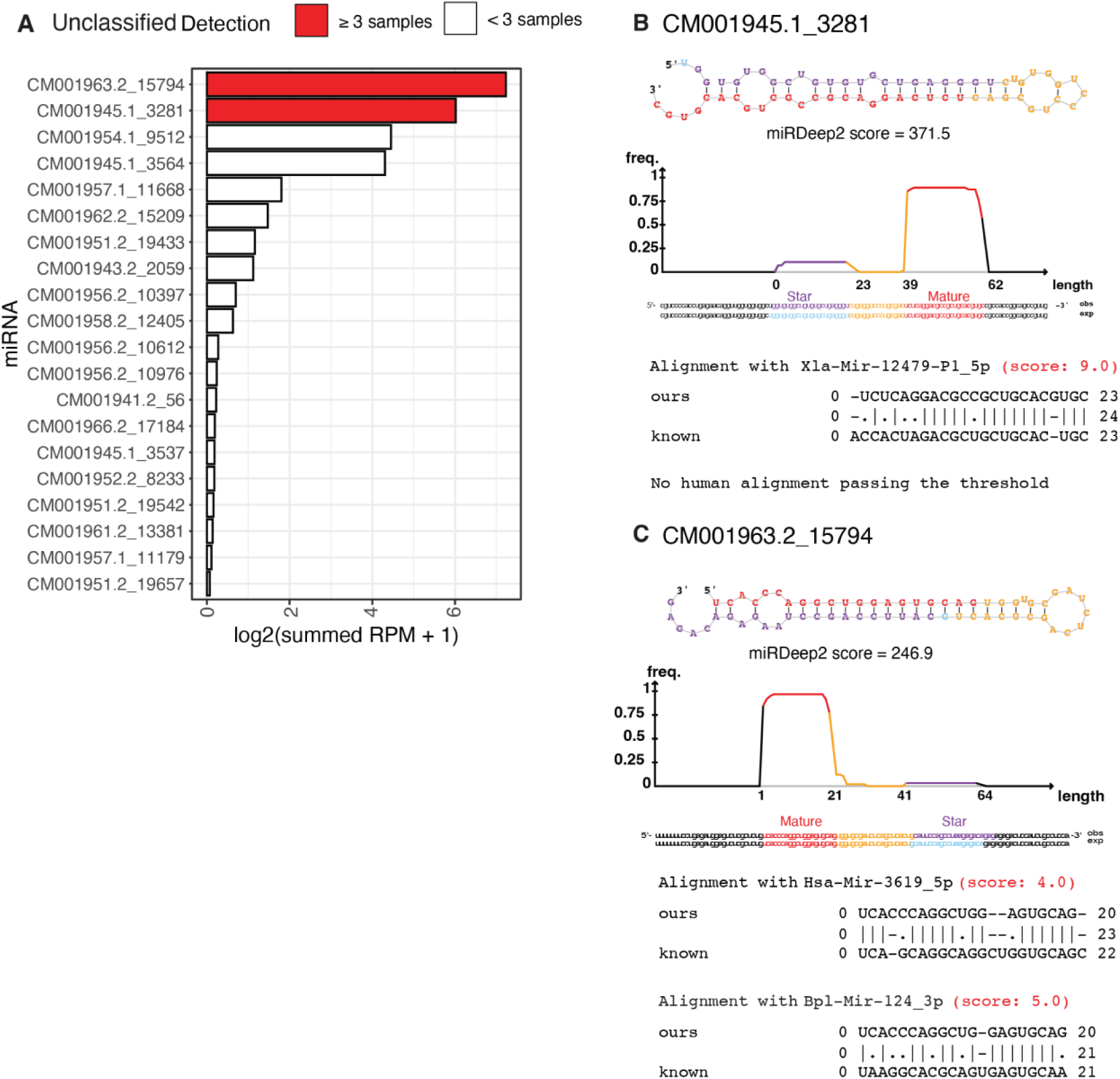
Unclassified miRNA candidates in African green monkey CSF miRNome. (A) Bar plot showing RPM for unclassified miRNAs with no family predicted by Rfam or MirMachine and alignment score < 34. Predicted miRNA sequences and alignment to mature miRNAs in MirGeneDB are shown for (B) CM001945.1_3281, and (C) CM001963.2_15794. In the precursor hairpins, red denotes the mature sequence, purple the star sequence, and yellow the loop sequence. Sequences in blue are those predicted by miRDeep2 based on the mature miRNA sequence present in the dataset. The alignments compare the dominant mature miRNA sequence in our dataset with the best-matched established mature miRNA sequences in MirGeneDB across all species, and the best matched human mature miRNA sequence, if an alignment exists that passes the threshold.

## DISCUSSION

Aging AGMs exhibit early-stage AD pathology without cognitive decline, while miRNAs can capture regulatory shifts that precede detectable changes in the proteome. Together, these offer complementary advantages for early-stage biomarker discovery that have not yet been leveraged in this model. Here, we take a first step by defining the AGM CSF miRNome antemortem. CSF is in direct contact with the central nervous system and provides a less invasive window into molecular processes in living animals. We designed a hierarchical annotation pipeline that first resolved family level conservation, then refined sequence level identity by alignment to known miRNAs. This approach allowed us to avoid missing conserved miRNAs that differed from reference sequences and prevented low confidence alignments to known miRNA families from being incorrectly classified as novel.

The AGM CSF miRNome is substantially conserved with humans. Of the 205 family-conserved miRNAs, 178 were canonical matches, 24 were conserved-3’-terminal variant, and three were noncanonical. 202 miRNAs were assigned to human reference sequences, supporting substantial conservation between AGM and human miRNAs. These miRNAs dominated the abundance distribution, and the ten most abundant were detected in every individual. A subset of these assignments was supported on an independent platform, with 150 human-annotated miRNAs also detected by the human-targeted qRT-PCR array and miRNA-seq abundance inversely correlated with Ct across overlapping miRNAs (Spearman ρ = −0.65, P < 0.001). The recovered miRNome were neurologically linked: 34 miRNAs showed preferential expression across brain regions in human tissue, predicted targets were enriched for axon guidance and synaptic organization, and disease enrichment spanned tauopathy and dementia. Alongside these, 20 candidates could not be assigned to any known family or sequence, two of which carry the read support and structural features expected of bona fide miRNAs. Together, these results establish a foundational AGM CSF miRNome for Alzheimer’s disease biomarker research and a reference for future comparative and translational work in this model.

AGM CSF miRNome includes miRNAs previously implicated in AD. Both family-conserved canonical and −3′-terminal variant miRNAs identified in AGM CSF have previously been implicated in AD. The family-conserved canonical miRNAs were associated with amyloid processing (hsa-miR-16-5p, hsa-miR-19b-3p, hsa-miR-29c-3p, and hsa-miR-125b-5p)^23–25^; tau-related kinase signaling (hsa-miR-125b-5p and hsa-miR-15a-5p)^26^; neuroinflammation (hsa-miR-146a-5p, hsa-miR-29c-3p, and let-7i-5p)^25,27,28^; synaptic function (hsa-miR-206 and hsa-miR-92a-3p)^29^; and mitochondrial or apoptotic regulation (hsa-miR-29b-3p and hsa-miR-204-5p)^30–32^. Notably, several 3′-terminal variant miRNAs also corresponded to miRNAs with established neurological or AD-related functions. These included individual members of the let-7 family, an evolutionarily conserved miRNA family associated with neuroinflammatory and neurodegenerative processes. The miRNA hsa-miR-124-3p is highly expressed in the brain and has been investigated as a potential AD biomarker, while experimental studies suggest that it may also have neuroprotective functions^33^. Additionally, hsa-miR-155-5p has been linked to aging, neuroinflammation, and AD-associated pathology^34^. Finally, hsa-miR-144-3p has been associated with AD, including through the regulation of pathways contributing to β-amyloid accumulation^35^. Consistent with this, predicted targets of the family-conserved canonical miRNAs were enriched for neurodegenerative disease categories, including dementia and tauopathy, further linking the AGM CSF miRNome to molecular processes implicated in AD. The recovery of these AD-associated miRNAs supports the translational potential of AGM CSF for human biomarker research.

Despite substantial conservation of the miRNome, a monkey is not a human. Sequence conservation therefore does not establish functional equivalence between species. Accordingly, the identification of AD-associated miRNAs linked to neuropathological processes should not be interpreted as evidence of functional equivalence between AGM and human. Even among conserved miRNAs, function also depends on expression level, tissue context, target availability, and physiological role. Conservation of the seed sequence may support overlapping target recognition even when miRNAs differ elsewhere in the mature sequence^36^, but it does not demonstrate equivalent function. Although CSF miRNAs remain relatively understudied in nonhuman primates, comparative brain studies have shown substantial conservation of miRNA composition across primates alongside species-specific expression patterns^37,38^. For example, 11% of miRNAs were differentially expressed in a human-chimpanzee prefrontal cortex comparison, compared with 31% in a human-rhesus macaque comparison^37,38^. This is consistent with greater divergence at greater evolutionary distance. Because rhesus macaques and AGMs are both Old World monkeys, the human-rhesus comparison likely provides a closer estimate of the expression differences expected between humans and AGMs than the human-chimpanzee comparison does. While this implies appreciable divergence, it nonetheless places monkey models closer to humans than rodent models. Determining whether the miRNAs identified here show comparable magnitudes and directions of expression in AGM and human CSF, particularly in the context of AD and related neuropathological diseases, will be an essential next step.

miRNA-seq and array data showed platform-specific biases when characterizing AGM miRNome. The qRT-PCR array detected a broader set of miRNAs than untargeted miRNA-seq. The array detected every miRBase annotated miRNA identified by miRNA-seq that was represented on the array, including a subset classified as family-conserved 3′-terminal variant from human reference sequences, as well as approximately 192 additional miRNAs that were not recovered by sequencing. The miRNAs detected only by the array generally had higher Ct values, consistent with the greater sensitivity of targeted amplification for detecting low-abundance miRNAs. Several brain-associated miRNAs detected by qRT-PCR were absent from the miRNA-seq dataset. These included miR-9-5p, a highly conserved miRNA with central nervous system functions across vertebrates^39^. Multiple members of the imprinted DLK1-DIO3 cluster were also detected, including hsa-miR-329-3p, hsa-miR-410-3p, hsa-miR-431-3p, and hsa-miR-433-3p^40^. In addition, the array detected the AD-related neuronal miRNAs hsa-miR-107^41^, hsa-miR-134-5p^42^, and hsa-miR-212-3p^43^. Their detection by qRT-PCR indicates that neurologically relevant miRNAs are present in AGM CSF but may fall below the detection limits of untargeted sequencing.

The family-conserved 3′-terminal variant miRNAs showed no significant correlation between sequencing and qRT-PCR, despite being detectable by both platforms. The human miRNA qRT-PCR assay could cross-detect closely related isomiRs that differ at their termini, potentially limiting reliable quantification of individual mature isoforms. This could produce two non-mutually exclusive sources of disagreement between platforms. First, miRNA-seq and qRT-PCR may not quantify equivalent miRNA species: sequencing resolves individual 3′ isomiRs, whereas qRT-PCR may capture signal from multiple closely related isoforms^44,45^. Second, sequence differences between an AGM isomiR and the canonical human target may influence amplification efficiency, resulting in unequal detection of different isoforms. Differences in isoform specificity and amplification efficiency may therefore contribute to the weaker cross-platform agreement observed for the family-conserved 3′-terminal variant miRNAs. However, this lack of significant correlation should also be interpreted cautiously given the relatively small number of these miRNAs detected across both platforms (n = 13), which limits statistical power to detect moderate cross-platform associations. Regardless of the source of the weaker cross-platform agreement, 3′ isomiR variation should be considered when developing miRNA assays for AGM translational studies, particularly when assays are designed using canonical human reference sequences.

Species-specific sequence variation and incomplete miRNA annotation make it difficult to determine whether an unclassified sequence is genuinely novel or simply absent from existing reference databases. Because AGMs are underrepresented in miRNA reference databases, the unclassified sequences identified here cannot yet be distinguished as putatively novel, lineage-specific, or conserved miRNAs that are absent from existing annotations. Two of the miRNAs were unclassified (CM001945.1_3281 and CM001963.2_15794) yet detected in four samples each and showed features suggestive of bona fide miRNAs. Both had a detectable star sequence (miRNA*) and high miRDeep2 scores, with strong read support across multiple libraries (Figure 5; Table S1). Mismatches and indels within and adjacent to the seed region indicate that these loci are sequence-distinct from their closest MirGeneDB matches and could alter target recognition, although their functional consequences remain to be validated, a pattern consistent with prior novel-miRNA reports^46,47^. Whether they are genuinely AGM-specific or simply absent from current references cannot be resolved from the present data, given how few non-targeted CSF miRNA studies exist and how sparse nonhuman primate data remain. One locus (CM001945.1_3281) matched the precursor of *Macaca fascicularis* novel_miR60-3p^48^, indicating the locus is at least conserved among nonhuman primates; but with no experimentally validated mature form in any species, its processing and relevance are undetermined. Targeted validation will be needed to place both loci.

Despite the absence of a previously defined AGM miRNome and a dedicated AGM miRNA reference annotation, our hierarchical annotation framework provides multiple lines of support for our assignments. Family- and sequence-level assignments were evaluated across broad cross-species references and then assessed separately against annotated human miRNAs, allowing us to distinguish deeply conserved candidates from 3′-terminal variant miRNAs without relying on a single annotation criterion. Following miRNA prediction, candidate annotation commonly relies on sequence similarity^46,47,49–51^. Our framework integrated sequence identity with family-level evolutionary conservation. Sensitivity analyses further showed that these classifications were stable despite the short length of miRNA sequences and the inherent limitations of alignment-based assignment. The convergence of these complementary approaches supports the robustness of our AGM CSF miRNome. This resource strengthens the molecular foundation for using AGMs as a translational model for miRNA-based biomarker discovery in early AD, while also identifying variation that may capture aspects of AGM biology not represented by human annotations.

The use of antemortem CSF prioritizes a valuable sample type that can be collected in vivo and repeatedly over the course of disease progression, which is particularly important in this early-stage model. Although CSF does not provide a direct representation of brain tissue, it offers a relevant window into molecular changes associated with the central nervous system and enables longitudinal biomarker assessment. However, CSF remains challenging to collect, and blood contamination can complicate interpretation of miRNAs^52^. Many miRNAs are detectable in both blood and CSF, including hsa-miR-16-5p, which was among the most abundant miRNAs in our samples^12^. Determining the extent to which such signals reflect CSF-resident versus blood-derived miRNAs will therefore be important. Defining which miRNAs are represented in CSF, how their abundance and isoform composition relate to direct measurements in brain tissue, and how these profiles change in disease will be important for interpreting their translational relevance. We have previously demonstrated proteomic shifts in CSF associated with this AD model^3^; an important next step will be to determine whether disease-associated changes are similarly detectable in the CSF miRNome.

The miRNome presented here was derived from female AGMs. This design reflects both the practical constraints and the translational strengths of the AGM model. Nonhuman primate studies are often limited by cohort size and sample availability^53^, and restricting this initial characterization to one sex reduced biological heterogeneity. Given the small sample size, we highlight miRNAs detected consistently in three or more individuals and indicate when specific miRNAs were detected in fewer individuals. The focus on females is also relevant because Alzheimer’s disease disproportionately affects women and sex differences may influence molecular pathways involved in neurodegeneration^54^. However, this design limits our ability to determine whether the AGM CSF miRNome differs by sex. Future studies should include larger cohorts of both male and female animals to define sex-specific and shared features of the AGM CSF miRNome.

By defining the AGM CSF miRNome, our study provides both a reference dataset and a practical framework for miRNA profiling in this longitudinal early-stage AD model. We demonstrate substantial conservation between AGM and human miRNomes while also highlighting the challenges of applying human-based annotation and detection approaches to a nonhuman primate biofluid. Our findings indicate that combining sequencing with targeted quantification will be important for characterizing miRNomes in AGMs and other understudied species. Future studies can build on this resource to determine how sequence conservation relates to miRNA abundance, isoform usage, and biomarker discovery across AGM and human systems. Moreover, given the limited characterization of CSF miRNomes in nonhuman species, this work also contributes to a broader understanding of species- and biofluid-specific variation in miRNA detection.

## METHODS

### AGM Study System and CSF sample collection

Six experimentally naïve female African green monkeys (*Chlorocebus sabaeus*) on the island of St. Kitts, West Indies, were used in this study. Animals were 7 to 9 years of age and did not have any clinical or pathological abnormalities. Age was estimated as previously described in Wakeman et al., 2022^55^. All *in vivo* procedures were performed by Virscio Inc. at their primate research facility located at the St. Kitts Biomedical Research Foundation (Lower Bourryeau Estate, St. Kitts and Nevis). Procedures were approved by the Virscio Institutional Animal Care and Use Committee (IACUC) and conducted in accordance with the Guide for the Care and Use of Laboratory Animals (National Research Council) and AAALAC International standards. CSF was collected into sterile tubes by cisterna magna tap. For our study, we used the CSF samples collected from AGMs at the baseline timepoint, following acclimation of the animals to the research environment.

### Total RNA extraction from whole CSF

CSF was thawed on ice, and then total RNA was extracted on the KingFisher™ Flex instrument system with the MagMAX™ mirVana™ Total RNA Isolation Kit (Thermo Fisher Scientific, Waltham, MA, USA), using the manufacturer’s protocol. 100 µL of whole CSF was treated with proteinase K at 65°C for 30 minutes. Following Proteinase K treatment, isopropanol, lysis buffer, and RNA isolation beads were added to the sample. As a spike-in control, 2.8 x 10^8^ copies of 5′-Phos-cel-miR-39-3p were added to the reaction. The reaction mixtures were processed on the KingFisher™ Flex instrument. The RNA concentration was measured using the Bioanalyzer 2100 (Agilent Technologies, Santa Clara, CA, USA). RNA extraction and quality control were performed at the Yale Center for Genome Analysis (YCGA).

### MiRNA library preparation and sequencing

5 µL of total RNA from each CSF sample was used as input in the QIAseq miRNA Library Kit (Qiagen, Venlo, Netherlands). Briefly, 3′ and 5′ adapter ligations were performed, followed by reverse transcription with primers containing unique molecular identifiers (UMIs). The cDNA was PCR-amplified to create the final library, which underwent a clean-up and quality control step prior to sequencing. The size and concentration of DNA in the final library was assessed using the Agilent TapeStation (Agilent Technologies, Santa Clara, CA, USA). The final quantification of the library was made using qRT-PCR using a KAPA Biosystems kit (Sigma-Aldrich, St. Louis, MO).

The concentration of samples was normalized to 2.0 nM, and the libraries were sequenced with 100 base pair paired-end sequencing on an Illumina NovaSeq instrument (Illumina, San Diego, CA, USA), according to the manufacturer’s protocol. Bacteriophage PhiX library (Illumina, San Diego, CA, USA) was added to the sequencing lane at 0.3% to verify the quality of the sequencing run. Raw data obtained from the sequencer was converted to base calls using the instrument’s Real Time Analysis (RTA) software. Library preparation and miRNA-seq were performed at YCGA.

### MiRNA sequencing data pre-processing

We assessed the quality of the sequencing fastq files using the FastQC module^56^. We processed fastq files with two rounds of fastp to remove the Illumina TruSeq 3′ and QIAseq miRNA adapters^57^. We then used these processed reads in all miRNA quantification and prediction programs discussed below. We performed fastp processing on a Linux Ubuntu virtual machine.

### Prediction of miRNAs

We aligned the processed reads to the *C. sabaeus* ChlSab1.1 GenBank assembly genome^58^ using mapper.pl, bypassing the adapter removal step -k <seq>^15^. In place of Bowtie v.1.1.1, we used Bowtie v.1.2.3 and the Intel Threading Building Blocks (*tbb*) library^59^. To assess the AGM as a model, we evaluated how its miRNome compares to that of humans. We characterized miRNAs by mapping the AGM miRNA-seq data to the human miRNA annotation database in MirGeneDB 3.0, which we chose over miRBase due to its lower false positive rate^18^. To identify miRNA candidates, we used miRDeep2.pl on a pooled fasta file containing collapsed reads from all samples, and human mature miRNAs as references, with no input reference files for *C. sabaeus* mature and precursor miRNAs step^15^. We then quantified miRNA read counts using miRDeep2 quantifier. Since no published annotations exist for *C. sabaeus* miRNAs, we used the output of miRDeep2 as custom reference files for quantifier.pl. We ran quantifier.pl with an added -k parameter to allow for different naming between the precursor and mature miRNAs. We executed all miRDeep2 scripts on a Linux virtual machine.

### Assignment of miRNAs to family-conserved and unclassified categories

As no miRNA reference exists for *C. sabaeus*, miRDeep2 cannot differentiate between known and novel miRNAs in the AGM. Therefore, we categorized miRNAs as family-conserved canonical, family-conserved 3′-terminal variants, family-conserved noncanonical and unclassified by examining whether they plausibly belong in established miRNA gene families or align to mature miRNA sequences in MirGeneDB.

We began by applying a threshold to the miRDeep2 output, retaining only miRNA candidates with a miRDeep2 score of ≥ 5 and total read count ≥10, to be conservative. We found that thresholds beyond 5 do not meaningfully reduce the number of retained candidates (Fig S1A). Additionally, we removed candidates for which an Rfam search identified them as non-miRNAs (details of this search are below). For each retained miRNA, we assessed whether it matched any known miRNA family using three complementary approaches (Fig S1B): MirMachine^16^, Rfam^17^, and sequence alignment to the MirGeneDB 3.0 database^18^.

For MirMachine, we used the precursor sequence with 40 base pairs of flanking sequence on both the 5′ and 3′ ends. The flanking sequences were obtained from the reference *C. sabaeus* genome sequence using the precursor start and end position reported by miRDeep2. Searches were conducted using default parameters against the Metazoa node, and only hits found on the same strand as the original miRDeep2 prediction were considered valid. The resulting family assignments were recorded under the column mm_family.

For Rfam searches, we used the cmscan tool from the Infernal suite with precursor sequences flanked by 40 base pairs on either side^17,60^. Searches were conducted using the flags -Z 0.227848, -cut_ga, --rfam, and --nohmmonly, following the Rfam documentation. The value of - Z is obtained by multiplying the length of the *C. sabaeus* genome by two, as instructed in the documentation. As with MirMachine, only strand-consistent miRNA hits were retained, and the family name was recorded under the column rfam_target_name.

Finally, we directly queried the mature miRNA sequence against MirGeneDB, identifying miRNAs by sequence alignment. Specifically, we conducted a global alignment between all pairs of candidates and mature sequences in the database. We used the Biopython v1.85 library to conduct the alignments with a match score of 2, mismatch score of −3, gap opening score of −5 and a gap extension score of −4. To determine an alignment threshold for a successful match, we leveraged alignments between MirGeneDB mature sequences. All pairs of alignments between mature sequences from the same family served as positive examples (n = 1,634,535 pairs), while alignments between sequences from different families served as negative examples. We randomly sampled n = 1,634,535 pairs of negative examples. An alignment score ≥ 0 corresponded to specificity of 99.7% and sensitivity of 83.3% (Figure S1D). Despite the lower sensitivity, all candidate miRNAs in our dataset had an alignment to a mature sequence that passed the threshold. All candidate miRNAs had a best alignment score ≥0; therefore, this threshold did not exclude any candidates from subsequent annotation. Furthermore, miRNA candidates were considered close mature-sequence matches when they had fewer than two mismatches, no more than five indels at the 3′ terminus, and no more than one indel outside the 3′ terminus. Final annotation categories were then assigned using both sequence-alignment characteristics and family-level support, as described below. All matched MirGeneDB IDs were reported as comma-separated values in the mirgenedb_ids_all column (Table S1).

To resolve redundancy, we deduplicated miRDeep2 output entries based on mature sequence identity. In cases where multiple entries shared the same mature sequence, we collapsed to the “family” level where no species is shown. Preference was given to entries with a MirMachine match, followed by those with a Rfam match, then a MirGeneDB match. MirMachine was prioritized since its training data is based on MirGeneDB^16^, which contains fewer false positives and false negatives than miRBase, while Rfam is trained on miRBase.

We first summarized the evidence supporting each provisional miRNA candidate across MirMachine, Rfam, and mature-sequence alignment using eight combinations (Table S1): MirMachine, Rfam, and high-alignment support (n = 156); MirMachine, Rfam, and low-alignment support (n = 21); MirMachine and high-alignment support (n = 2); MirMachine and low-alignment support (n = 3); Rfam and high-alignment support (n = 22); Rfam and low-alignment support (n = 7); high-alignment support only (n = 2); and low-alignment support only (n = 35).

Because multiple provisional candidates could converge on the same miRNA assignment across the alignment, MirMachine, and Rfam analyses, we collapsed these entries to a single record per miRNA to avoid double counting. When provisional candidates mapping to the same miRNA received different annotations, we retained the highest-ranking assignment. Following this consolidation, miRNAs were assigned to four final annotation categories. Family-conserved canonical miRNAs (178) were defined by complete mature-sequence identity to an established MirGeneDB reference. These exact matches also exceeded the alignment-score threshold of ≥34. Family-conserved 3′-terminal variants (n = 24) had family-level support from MirMachine and/or Rfam, identical 5′ and seed sequences to the assigned reference, no internal sequence differences, and variation confined to the 3′ terminus. Family-conserved noncanonical miRNAs (n = 3) retained family-level support from MirMachine and/or Rfam but showed seed or internal sequence differences, or other sequence divergence that could not be explained solely by 3′-terminal variation. Candidates with no family assignment from MirMachine or Rfam and no close mature-sequence match to an established MirGeneDB reference were classified as unclassified (n = 20). Together, these categories comprised 205 family-assigned miRNAs and 20 unclassified candidates. Precursors of unclassified miRNAs detected in three or more samples were additionally queried against the NCBI core nucleotide database using NCBI BLASTN (megablast)^22,61^.

### Functional Enrichment and Tissue Expression Analyses

To investigate the relevance of miRNAs to cellular function, we performed target prediction and enrichment analysis using a list of miRNAs satisfying two conditions: (1) having an alignment score greater than or equal to 34 mapped to a MirGeneDB human miRNA reference, and (2) having a family detected by one or more of the following methods: Rfam and MirMachine (mirgenedb_ids_hsa - Table S1). We used the human genomic coordinates file downloaded from MirGeneDB 3.0 to convert MirGeneDB IDs to miRBase accession numbers^16^. These accession numbers were then used as input for target prediction for each miRNA using the miRWalk platform, which integrates results from multiple target prediction tools^21^. We retained only those target genes that were predicted by both TargetScan and miRDB to increase confidence in target assignments.

We conducted a functional enrichment analysis of miRNA target genes using the R package clusterProfiler^62^. We carried out an Over-Representation Analysis (ORA) to identify significantly enriched biological pathways and ontologies among the predicted gene targets. Specifically, we assessed enrichment across the Kyoto Encyclopedia of Genes and Genomes (KEGG), Gene Ontology (GO), Disease Ontology (DO), and Reactome pathway databases^63,64^. We implemented Disease Ontology analysis through the DOSE package^65^, and Reactome pathway enrichment using the ReactomePA package^66^. We conducted all analyses in R and used default parameters unless otherwise specified.

To assess the tissue distribution of human homologs corresponding to AGM CSF miRNAs, we examined their expression across human tissues using GTEx v10^20^. For miRNA expression, we analyzed family-conserved canonical miRNAs and miRNAs detected only by the array using the GTEx small RNA-seq dataset. We used the normalized miRNA TPM matrix and calculated the median expression of each miRNA within each tissue across GTEx samples. We then assessed whether the miRNA had preferential expression in specific organs or tissues.

### Profiling of miRNAs with the ID3EAL^TM^ PanoramiR panel

MiRNA profiling was performed by the Detection Unit, Precision RNA Medicine Core at Beth Israel Deaconess Medical Center (RRID: SCR_024819) using the quantitative RT-PCR-based ID3EAL™ PanoramiR miRNA Knowledge Panel (Mirxes, Singapore)^19^, which quantifies 376 human miRNAs per sample. 20 µL of CSF total RNA per sample was reverse transcribed according to the manufacturer’s protocol, at 25°C for 10 min, 30°C for 10 min, 35°C for 10 min, 40°C for 10 min, followed by RNA degradation at 95°C for 5 min.

The resulting cDNA was pre-amplified at 95°C for 10 min and 40°C for 5 min, and 8 cycles of 95°C for 10 s and 60°C for 30 s. Pre-amplified cDNA was used in the following qRT-PCR cycling protocol: 1 cycle of 95°C for 10 min, and 40 cycles of 95°C for 10 s and 60°C for 40 s, using the Quant Studio 5 RT-PCR system (Thermo Fisher Scientific, Waltham, MA, USA).

We processed raw cycle threshold (Ct) values in R. We assigned a Ct value of 40 to undetectable miRNAs. We corrected for intra-run variability by using the Ct values of the RNA spike-in controls. To perform global normalization, we imputed missing data points using the mean ± 3 standard deviations for each miRNA.

### Correlation Analyses with Sequencing Abundance and Ct values

To evaluate consistency between miRNA-seq and qRT-PCR-based measurements and to provide orthogonal support to our sequence derived miRNome, we assessed the relationship between miRNA-seq read counts for AGM miRNAs assigned to human references and qRT-PCR Ct values.

We converted MirGeneDB identifiers from miRNA-seq data to miRBase nomenclature for comparison with the PanoramiR array. Assignments that could be mapped unambiguously to a single miRBase target were retained, including manually resolved one-to-one mappings. Assignments corresponding to multiple possible miRBase paralogs were excluded from the one-to-one cross-platform comparison because the specific array target could not be uniquely identified. These unresolved assignments included Hsa-Mir-135_5p, Hsa-Mir-181_5p, Hsa-Mir- 320_3p, and Hsa-Mir-376_3p. After mapping, we calculated Spearman correlations between mean reads per million (RPM) from sequencing and mean Ct values for each miRNA detected on both platforms.

## SUPPLEMENTARY TABLES

**Table S1. miRDeep2-predicted miRNA candidates and hierarchical annotation results.** For each provisional miRNA candidate, the table reports the miRDeep2 prediction score; read support for the predicted precursor, mature, loop, and star sequences; randfold significance; and any Rfam alerts. Hierarchical annotation results include the assigned annotation category, miRNA family, Rfam target, matches to miRBase and MirGeneDB, and whether the candidate met alignment-score criteria across the full reference database and against annotated human miRNAs. An example miRBase miRNA sharing the same seed sequence is provided where available. Consensus mature, star, and precursor sequences and the genomic coordinates of each predicted precursor are also reported.

**Table S2. miRNA detection by the PanoramiR array and overlap with miRNA sequencing.** The full PanoramiR miRNA array results are provided, including the mean cycle threshold (Ct) value for each miRNA across samples. miRNAs are further classified according to whether they were detected by both the PanoramiR array and miRNA sequencing, uniquely by the array, or uniquely by sequencing. The table also provides the conversion between MirGeneDB identifiers and miRBase names.

**Table S3.** GTEx tissue-expression profiles of human miRNA homologs corresponding to AGM CSF miRNAs. The table includes GTEx expression data used to evaluate the tissue-expression patterns of human miRNA homologs corresponding to miRNAs detected in the AGM CSF miRNA-seq dataset.

**Table S4. MirGeneDB accession numbers and predicted mRNA targets of miRNAs detected by miRNA sequencing.** The table includes accession numbers corresponding to the assigned MirGeneDB miRNA identifiers and miRWalk-predicted mRNA targets for miRNAs detected in the miRNA-seq dataset.

**Table S5. Functional enrichment analysis of miRNAs detected by miRNA sequencing.** The complete functional enrichment output is provided for miRNAs detected in the miRNA-seq dataset, including enriched pathways, biological processes, and disease-associated terms.

## SUPPLEMENTARY FIGURES

**Figure S1.**
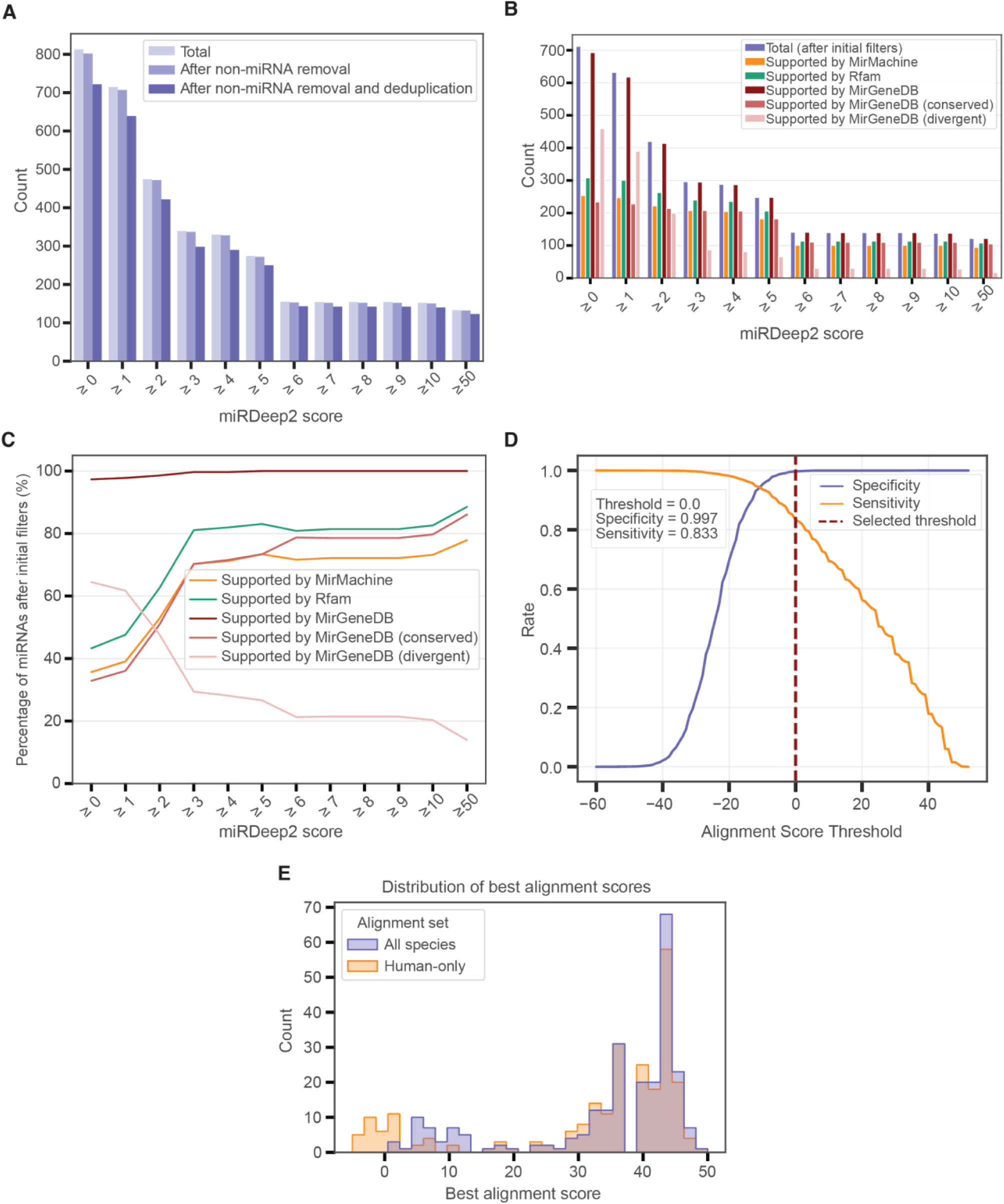
Filtering miRNAs and determining alignment score thresholds. (A) miRNAs from the miRDeep2 output were deduplicated and filtered to retain candidates with a miRDeep2 score ≥ 5, and sequences mapping to non-miRNA small RNAs in Rfam were removed. (B-C) Retained miRNAs were assessed to determine whether they plausibly belonged to established miRNA families based on Rfam and MirMachine assignments or sequence similarity to miRNAs in MirGeneDB 3.0. (D) An alignment score threshold of ≥ 0 had a 99.7% specificity and 83.3% sensitivity. A more stringent threshold of ≥ 34 (99.7% specificity and 40.0% sensitivity) was used to distinguish family-conserved canonical, family-conserved 3’-terminal variant, family- conserved noncanonical, and unclassified. (E) Distribution of best alignment scores for alignments against miRNAs across all species represented in MirGeneDB (purple) and against human miRNAs in MirGeneDB (orange).

**Figure S2.**
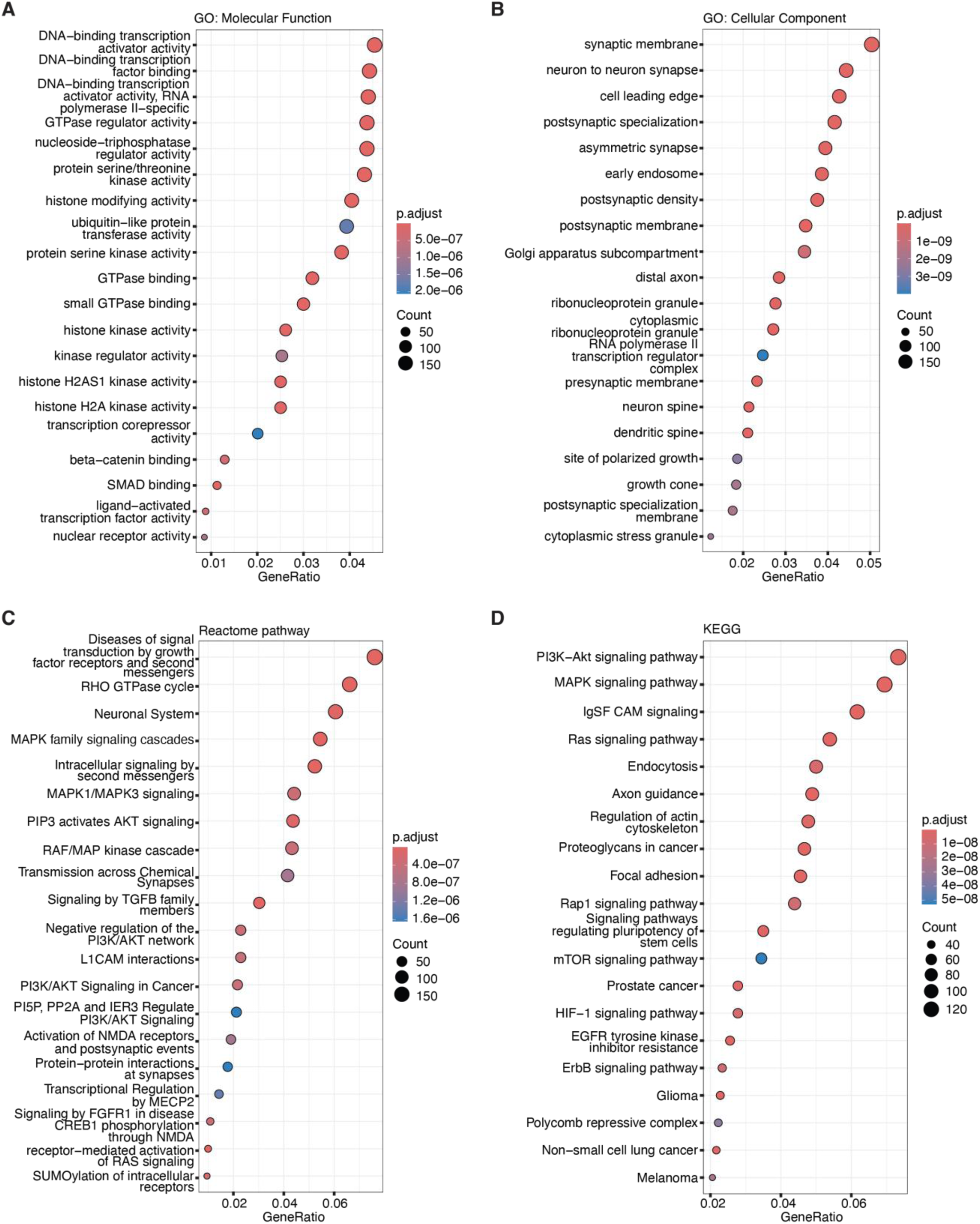
Gene ontology (GO), reactome, and KEGG pathway enrichment of mRNA targets from the miRNA-seq dataset. The top 20 enriched GO molecular function (A), cellular component (B), reactome pathway (C), or KEGG pathway (D) terms for mRNAs predicted to be targets of family-conserved canonical miRNAs (≥ 34 alignment score with human reference, with a family detected by MirMachine and/or Rfam) by miRWalk, miRDB, and TargetScan. The GeneRatio indicates the number of mRNA targets to the total number of mRNA genes in the input list, while the terms are ranked by adjusted p-values.

**Figure S3.**
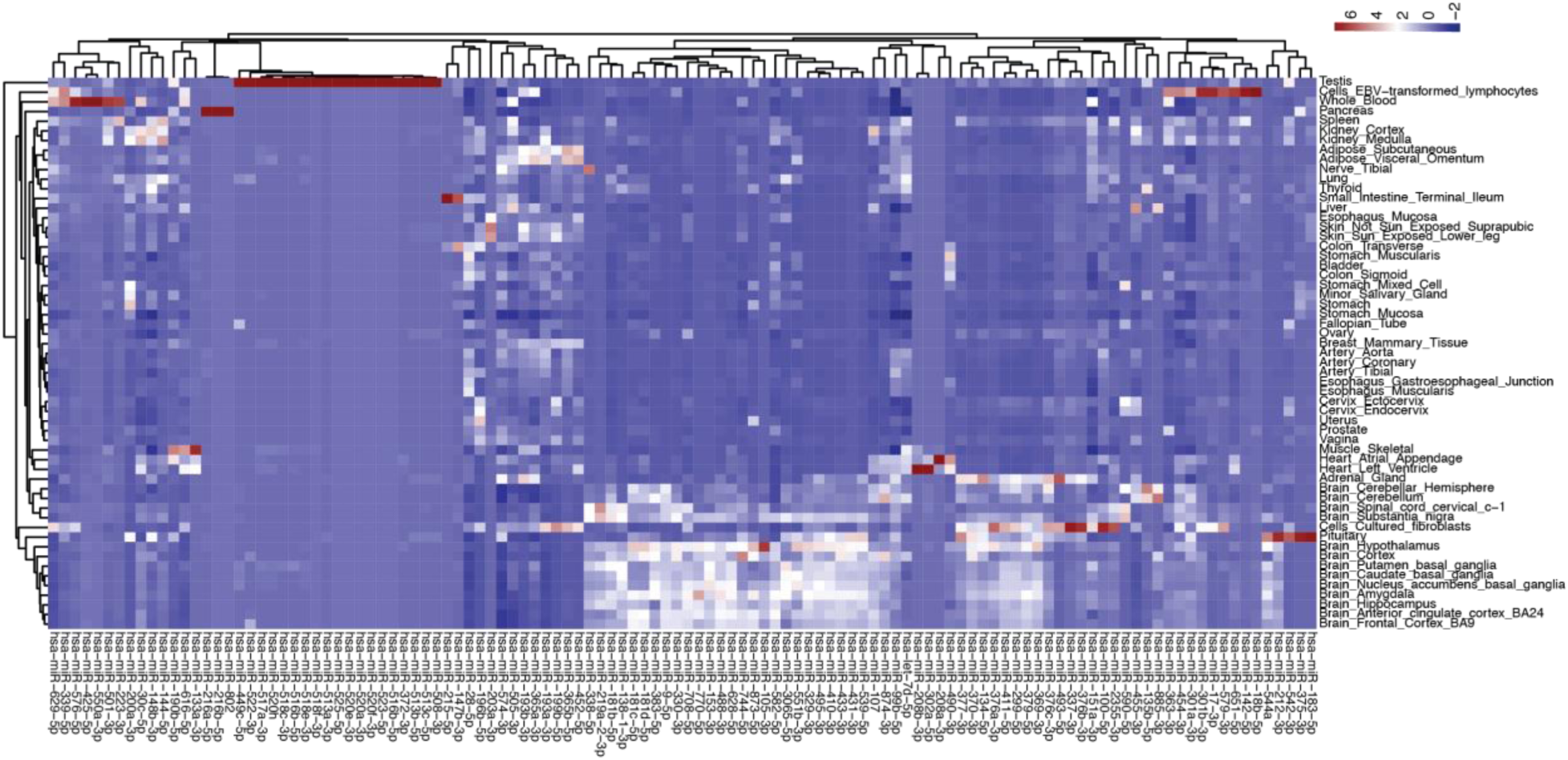
Tissue expression analysis of miRNAs unique to qRT-PCR array dataset. Tissue expression analysis of miRNAs unique to the PanoramiR qRT-PCR array dataset, using GTEx v 10.

## AUTHORS CONTRIBUTION

**Conception:** SD, BRPB

**AGM experiment and sample collection:** MRW, JDE, MSL

**Methodology:** SD, RA, XL, CSB, ACR, PNY, WAH, FJS, ADM, BRPB

**Data curation:** SD, RA, XL, YJH, BRPB

**Formal Analyses:** SD, RA, XL, GG, BRPB

**Visualization:** SD, RA, XL, BRPB

**Writing - initial:** SD, BRPB

**Writing - review & editing:** All authors

**Software:** SD, RA, XL, BRPB

**Supervision:** BRPB

**Project administration:** BPRB

**Funding acquisition:** ADM, MRW, GG, BRPB

## ACKNOWLEDGEMENTS

We thank the Yale Center for Genome Analysis, with support from NIH grant 1S10OD030363- 01A1, for assistance with RNA isolation, library preparation, and sequencing, and the Detection Unit within the Precision RNA Medicine Core at Beth Israel Deaconess Medical Center, Harvard Medical School (RRID: SCR_024819), for running the PanoramiR arrays.

## FUNDING INFORMATION

This work was supported by the National Science Foundation Postdoctoral Fellowship awarded to BRPB, the Small Business Innovation Research grant AG067832 awarded to MW, the National Institute of Health Office of Director grant R03OD036491 awarded to GG, the National Institute on Aging (NIA) grants R01AG058816 and R01AG082093 to FJS, and NIA grant R01AG068285 awarded to ADM.

## DATA AND AVAILABILITY

- The raw miRNA sequence FASTQ files is deposited in the NCBI Sequence Read Archive under BioProject PRJNA1503537. Reviewer link: https://dataview.ncbi.nlm.nih.gov/object/PRJNA1503537?reviewer=bsrsrkvq8ft26hof7t12cl1krr
- Code for data processing and analyses is available at: https://github.com/Biancabrown/African-Green-Monkey-MiRnome

## RESOURCE AVAILABILITY

Requests for additional information and resources should be directed to and will be fulfilled by the lead contact, Bianca Brown.

## Notes

https://github.com/Biancabrown/African-Green-Monkey-MiRnome

### Competing Interest Statement

The authors have declared no competing interest.

